# The impact of musical expertise on disentangled and contextual neural encoding of music revealed by generative music models

**DOI:** 10.1101/2024.12.20.629729

**Authors:** Gavin Mischler, Yinghao Aaron Li, Stephan Bickel, Ashesh D. Mehta, Nima Mesgarani

**Affiliations:** Mortimer B. Zuckerman Mind Brain Behavior Institute, Columbia University, New York, NY 10027, USA; The Feinstein Institutes for Medical Research, Northwell Health, Manhasset, NY, 11030, USA

## Abstract

Music perception involves the intricate processing of individual notes and their contextual relationships within a piece. However, how the brain encodes and organizes these features, particularly in relation to musical expertise, remains unclear. Using noninvasive and invasive electrophysiological recordings alongside generative music models, we reveal that musicians exhibit neural encoding that is more attuned to the separation and integration of complex musical structures, with a pronounced left-hemispheric bias compared to non-musicians. Invasive recordings further highlight a hierarchical and spatially organized representation of musical features and context across brain regions. These findings advance our understanding of how the brain processes music and the role of musical training in shaping auditory cognition.

## Introduction

Music perception represents a remarkable feat of cognitive processing. With its intricate tapestry of pitch, rhythm, and harmony, music is a unique and complex auditory stimulus that engages a wide variety of neural processes (*1–7*). The brain not only deciphers individual notes, characterized by features such as pitch, duration, and timbre, but also integrates these elements into rhythmic patterns, melodic contours, and larger musical sequences. This process spans multiple timescales, from the rapid recognition of individual notes and intervals to the apprehension of melodic contours and rhythms, and up to the integration of musical phrases and thematic structures (*8–11*). The perception of music and the pleasure derived from it are intimately connected to the predictability and surprisal of musical events (*12–15*). Hence, the ability to integrate and interpret contextual cues is pivotal for the appreciation and understanding of music, which is also shaped by musical training (*16*). The auditory cortex’s role in music perception is further highlighted by its selectivity for musical features such as pitch (*6*, *17*, *18*), rhythm (*19*, *20*), and timbre (*17*, *21*). Higher order areas of the auditory pathway, and in particular the inferior frontal gyrus, exhibit a complex, hierarchical neural encoding adept at both fine-grained acoustic analysis and higher-level structural and syntactic awareness (*22–26*). The neural responses to musical notes also encode their predictability (*13*, *27*), a feature that relies on contextual processing of higher-order structure. The precise nature of this context-dependent neural encoding across various timescales and its modulation by musical expertise remains unknown. Musicians develop heightened automatic auditory sensitivity to melodic contours (*28*) and enhanced predictive capabilities (*29*), leading to greater perception and enjoyment of complex musical compositions (*16*). Furthermore, musicians may recruit distinct neural regions, including increased involvement of the left hemisphere, compared to non-musicians, during music note and low-level musical feature analysis (*30–32*). This suggests that the musician’s brain constructs an enhanced representation of music, facilitating heightened sensitivity to musical features and improved predictive capabilities. However, the neural basis of this enhanced representation, including how hemispheric recruitment patterns support the hierarchical processing of musical features remains unclear.

A major challenge in studying the cortical encoding of musical context is the difficulty in measuring and quantifying context in music at various timescales. Previous studies approached this problem with a scrambling paradigm, showing that degradation of the musical structures at different timescales, such as randomizing the pitch or timing of notes, results in reduced neural encoding in different brain areas (*3*, *22*, *33*). Other studies used computational models with limited temporal dependencies to measure the predictability of musical notes (*13*). Past studies of deep learning model features in musical encoding have shown that they can be used to estimate the melodic expectation and surprise in listeners (*27*, *34*), though they did not investigate the properties of the representations or their relation to musical training. The advent of computational models capable of naturalistic music generation (*35*, *36*), particularly transformer models, presents an unprecedented opportunity to probe the depths of contextual processing in the human auditory system during music perception. These models, capable of handling sequential data over multiple timescales and mimicking the hierarchical nature of musical structure, offer a parallel for examining how the brain might organize and interpret musical information. By simulating the complex layering of musical context, these models enable a comparative analysis between the model and the brain that can advance our understanding of the capacity of the auditory processing pathway for contextual integration and how it changes with musical expertise.

In this study, we investigate how the human auditory pathway encodes the structure and context of musical stimuli, and how this encoding is influenced by musical training. We used electrophysiological recordings from both the scalp (EEG) and depth and surface intracranial electrodes (iEEG) as subjects listened to a series of musical compositions. Linear predictions of the neural responses from various layers of a transformer model trained to produce music revealed the encoding of hierarchical and context-dependent musical features in the brain.

Furthermore, the pattern of disentangled feature encoding showed significant distinctions between musicians and non-musicians. In addition, the intracranial recordings illuminated spatial variations of this encoding, elucidating the role of training and exposure in shaping the cortical representation of music and offering insights into the brain regions responsible for these neural representations.

## Results

To examine the cortical encoding of musical context, we recorded scalp EEG from 10 expert pianists with a degree in music and a minimum of ten years of music training (musicians) and 10 subjects without musical training (non-musicians) (Fig. 1a) (*13*). We also recorded intracranial EEG (iEEG) from 6 neurological patients undergoing epilepsy surgery, none of which had any musical training. All subjects listened to 30 minutes of music consisting of 8 Bach piano pieces. We used a 13-layer sequence-to-sequence transformer model to model the same musical pieces (Musicautobot (*37*)). The model was trained on music MIDI data, including classical music from a wide range of composers, and consisted of both an encoder and a decoder, and was trained on multiple tasks including masked note, chord, and melody prediction. We derived causal transformer features from all layers of the encoder to understand the representations learned by the model for processing continuous music, which we then compared to the neural recordings from subjects listening to the same piano pieces.

**Fig. 1.**
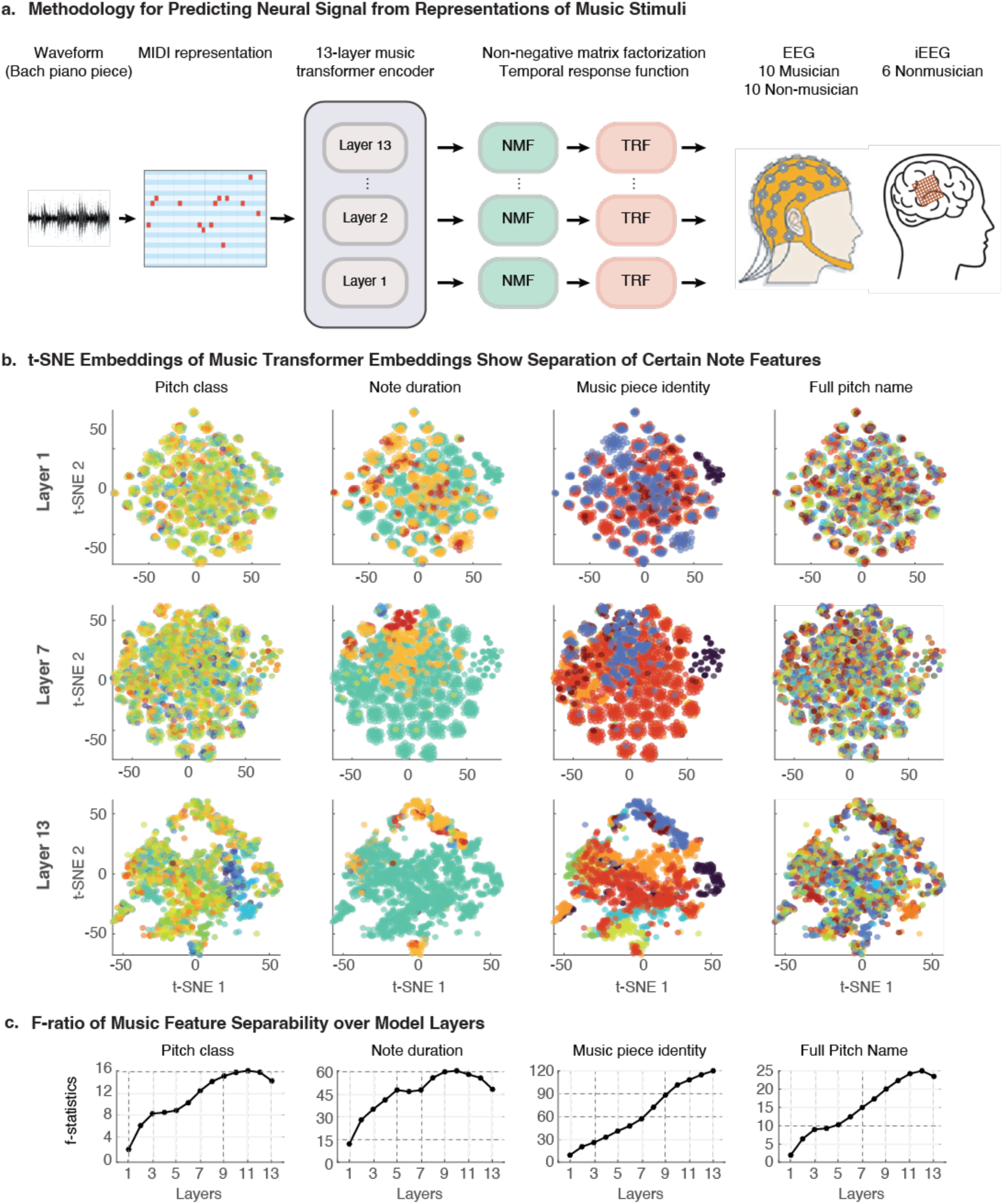
Investigating music transformer embeddings. **a)** The paradigm for predicting neural data from a generative music transformer model. Bach music pieces were fed into a pre-trained encoder-decoder transformer model using their MIDI representation, and then the note embeddings from each of the initial embedding layer and subsequent 12 layers of the encoder were extracted for further analysis. The dimensionality of these embeddings was reduced via nonnegative matrix factorization (NMF), and then cross-validated temporal response function (TRF) models were trained to predict the EEG and iEEG electrode responses. **b)** Note embeddings from layers 1, 7, and 13 of the transformer model were plotted in two dimensions with t-SNE, and notes were colored based on either their pitch class, note duration, music piece identity, and full pitch name. **c)** To measure the separability of each illustrated feature from the embedding space, the F-ratio from the full-dimensional embeddings was computed from each layer of the model.

### Increased contextual encoding in transformer layers

We first quantified the encoding of music features in transformer layers. These models have been shown to generate contextualized embeddings reflecting not only the note itself, but its relationship with other notes. The architecture of transformer models in language processing is characterized by a progression from encoding simple, low-level information in the initial layers to capturing more detailed, higher-order features in the deeper layers (*38*). We examined this gradual encoding in the music transformer model to understand whether it demonstrated a similar hierarchical encoding of features based on their complexity and departure from simple acoustic features. First, we projected the embeddings from each layer to two dimensions using t-SNE (*39*). Figure 1b illustrates two-dimensional t-SNE features for layers 1, 7, and 13 of the transformer models. In Figure 1b, data points represent individual music notes across all music pieces and are colored based on a certain music feature, including pitch class (chromatic categories, e.g., C, C#), note duration (normalized temporal lengths), music piece (assigned from MIDI metadata), and full pitch name (combination of pitch class and octave, e.g., C4, G#3). As we progress from layer 1 to layer 13, clear clusters for note duration begin to form by layer 7 and become more distinct by layer 13, indicating that note duration is encoded more explicitly in deeper layers. In contrast, more abstract attributes, such as the identity of the music piece to which a given note belongs, requires a broader understanding of musical context and structure. t-SNE embeddings do not show obvious clustering based on this attribute at layer 7 (Fig. 1b).

However, by layer 13, more distinct clusters emerge, demonstrating that the deeper layers of the transformer model contain more disentangled representations of these features. Further evidence of this progressively hierarchical encoding comes by computing the f-ratio of these discrete note classes from the embeddings, a measure of the separability of the classes in the high-dimensional space. We measured the f-ratio between notes grouped by either pitch classes, note duration, music piece, and full pitch names using the full representations from layers 1 to 13. The f-ratio between these note features generally increases over layers (Fig. 1c), indicating that the representations become increasingly separable over layers based on these attributes. These findings confirm that the music transformer model’s embeddings are extracting relevant features from the music pieces which go beyond simple acoustic features, and that later layers more robustly separate various characteristics of the notes that likely require integration over longer timescales and greater contextual information.

### Progressive neural encoding of transformer music features

To understand the extent to which the cortical surface responses to the music reflect the embeddings of the transformer models, we predicted the EEG response of each electrode for all 20 subjects from the transformer features. This was done after applying nonnegative matrix factorization (NMF) for dimensionality reduction. These transformed features were then used to linearly predict the EEG data using temporal response functions (TRF) with leave-one-out cross-validation for each subject and music piece (*40*, *41*). We analyzed the correlation between actual and predicted neural responses with cross-validation. We used Pearson correlation to assess the prediction performance with cross-validation (see Methods). Given that the first layer embeddings provide the least contextual and separable feature encoding, we computed the improvement in prediction correlation by each subsequent layer compared to the first layer.

Figure 2a shows these correlation improvements over the layers of the transformer, averaged over the musician and non-musician groups. Both populations showed an increase in prediction correlations over layers, meaning that the more disentangled note representations formed in the later layers were more predictive of the EEG responses. This suggests that across subject groups, the encoding of musical features in the brain is a progressive process that unfolds over multiple levels of neural processing, with separable representations of musical features arising in both musicians and non-musicians.

**Fig. 2.**
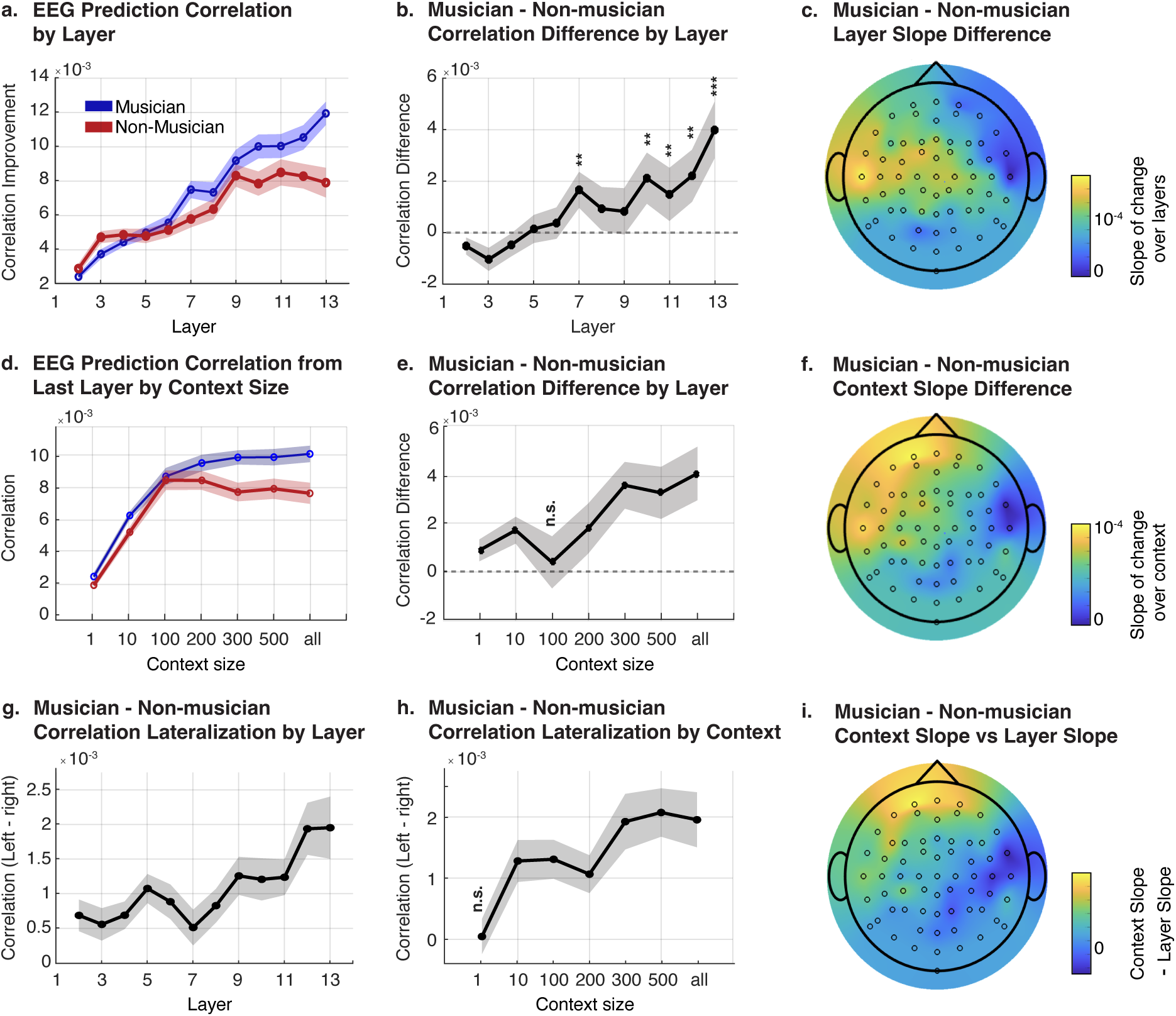
Results from the analysis of EEG recordings from musicians and non-musicians. **a)** EEG prediction correlation improvements compared to the first layer (averaged over all electrodes from each subject) were grouped into musician and non-musician brains and averaged. The plot shows the average over subjects at each transformer layer, and shaded regions indicate standard error of the mean over subjects. **b)** The difference between musician and non-musician EEG prediction correlations is shown over model layers. The significance of this difference was computed with a t-test, and with statistical significance indicated by stars above each significant layer (** indicates p<0.01, *** indicates p<0.001). The shaded region indicates the average standard error of the musician and non-musician sub-groups. **c)** The slope of the correlations over layers was computed by fitting a line on each electrode’s correlations. The slopes were averaged for each electrode across musicians and non-musicians separately, then the difference between the average slopes for these groups was plotted on a scalp topoplot, highlighting the more positive slopes of musicians especially in the left hemisphere. **d)** Average EEG prediction correlations using the final layer’s embeddings are shown for a range of limited context sizes, where the model was only given a certain number of previous notes as input. Shaded regions indicate standard error of the mean. **e)** The difference between musician and non-musician EEG prediction correlations is shown over context sizes. The significance of this difference was computed with a t-test, and all context sizes were statistically significant except for 100 notes. **f)** The slope of the correlations over context sizes was computed by fitting a line on each electrode’s correlations. The slopes were averaged for each electrode across musicians and non-musicians separately, then the difference between the average slopes for these groups was plotted on a scalp topoplot, illustrating the more positive slopes of musicians especially in the left hemisphere and frontal region. **g)** Lateralization of EEG prediction correlations over layers was computed by averaging the prediction correlations over the left and right hemisphere electrodes. Then the difference between the average musician and average non-musician lateralization was plotted, with the shaded region showing the average standard error of the mean between musicians and non-musicians. A t-test for statistical significance of the difference between musicians and non-musicians found every layer to be significantly different (p<0.05). **h)** Lateralization of the EEG prediction correlations from the final layer’s embeddings was computed for different limited context sizes, and the difference between the average musician and non-musician lateralization was plotted. Shaded regions indicate the average standard error of the mean between musicians and non-musicians. A t-test for statistical significance of the difference between musicians and non-musicians found every context size except for 1 note to be significantly different (p<0.05). **i)** The additional slope difference from context slope compared to layer slope differences between musicians and non-musicians (i.e. Fig 2c subtracted from Fig 2f).

### Differential neural encoding of music from expertise

We then explored the impact of musical expertise on the strength of this encoding of disentangled musical features by directly comparing the musician and non-musician groups.

Although the correlation generally increased over layers for both groups, the average correlation is significantly higher for musicians than non-musicians in nearly all the later layers of the model (Fig. 2b). In fact, the correlation values plateaued for non-musicians in the last four layers of the model, while they continued to increase for the musician group, leading to the difference in correlation between musicians and non-musicians generally increasing over transformer layers (Fig. 2b). This finding suggests that musical expertise facilitates the formation of more disentangled musical features which are more similar to those created by the later layers of a transformer model.

Since layer progression is related to contextual encoding in transformer language models (*42*, *43*), we hypothesized that contextual encoding may play a significant role in the observed patterns in the music transformer. To test this hypothesis, we predicted the EEG responses from the final transformer encoder layer as we varied the contextual input size to the model. The context size refers to the number of music notes fed into the model before the current note, thus limiting the maximum information available to the network. We varied the input size from 1 (a single note with no context) to full context (in which all notes before the current note were included in the input). For musicians, we found that the final layer EEG prediction correlations continued to increase as the context size grew larger, even until an input size of 300 notes (Fig. 2d). Non-musicians, in contrast, did not show improvement after an input size of 100 notes. The difference between musicians and non-musicians over context also continues increasing over layers (Fig. 2e), and musicians always showed significantly higher correlations than non-musicians except at a context size of 100 notes (t-test, p < 0.05). These findings suggest that musicians integrate musical context over longer time periods, while non-musicians may be limited in integrating extended musical contexts, reaching a plateau beyond a certain contextual window.

This could be attributed to the musicians’ enhanced cognitive functions and neural mechanisms for processing complex musical structures (*44*), including their ability to anticipate and integrate long-term dependencies in music (*45*). Our findings demonstrate that strongly disentangled musical features and contextual information are more strongly encoded in musicians, underscoring the profound impact of musical training on the brain’s capacity to process and understand music and providing further evidence for the plasticity of the human auditory system in response to specialized training.

### Spatial variations in music context encoding and hemispheric lateralization of musical experience

We systematically quantified the effects of contextual encoding of each electrode by measuring its slope of change in the prediction correlation (y) as a function of either transformer layers or context window (x) to calculate the slope of change (a) in the equation y = ax + b. A higher coefficient indicates a larger improvement over the baseline (first layer or no context) as either the layers or context window increases, and therefore it suggests that more separable features or extended context are more important for that electrode when processing the music stimuli.

We visualized the spatial variation of slopes over layers and context windows on topographic maps, focusing on the difference between musicians and non-musicians. This uncovered hemispheric differences between these subject groups, where the prediction correlations of the left hemisphere in musicians displayed a more positive correspondence with model layer and context length than in non-musicians (Fig. 2c and 2f). Also, the context length had a larger impact on both the left temporal and frontal regions, whereas the layer effect was more localized to the left temporal region (Fig. 2i), suggesting recruitment of slightly different cortical regions for processing disentangled and contextual music features.

Given the left-hemispheric lateralization we identified in the slopes of prediction correlations with layers and context lengths, we summarized the impact of musical training on the lateralization of EEG prediction correlations in further analyses. For each subject, we estimated the left hemispheric lateralization by computing the difference in prediction correlation between the left and right hemisphere electrodes. Then we computed the difference in this lateralization value between the musician and non-musician at each layer of the model. We found that musicians showed significantly higher left hemisphere lateralization in all layers of the model compared to non-musicians (t-test, p < 0.05) (Fig. 2g), and this difference became even more pronounced in later layers. Similarly, by varying the context size and using the correlations from the last layer only, we found that musicians had significantly higher left hemisphere lateralization at all but a single note of context (t-test, p < 0.05) (Fig. 2h). These findings suggest that the left hemisphere may be more involved in the disentanglement of musical features and that frontal areas may play a more significant role in the increased encoding of musical context after musical training. These findings highlight the differences in where and how disentangled musical features and long-term contextual information emerge with musical training.

### Intracranial EEG analysis of brain regions involved in contextual music encoding

While EEG is a method that allows for a better control of subjects (e.g. musicians vs. nonmusicians), it does not have enough spatial resolution to examine the brain regions responsible for different contextual encoding. Therefore, we extended our analysis to intracranial EEG (iEEG) from 6 subjects, using a similar analysis approach as the one used for EEG. Although these human subjects were not trained musicians, the anatomical recording specificity allowed for new insights into the locus of music-transformer brain similarity. The iEEG signals were predicted using transformer features across layers in the same way as was done with EEG recordings, and we identified a rise in cross-validated prediction correlation over layers for most electrodes, though not all. We performed unsupervised k-means clustering on the normalized correlations over layers, identifying three clusters of electrodes (Fig. 3a). These clusters significantly differed in which layers scored highest, as quantified by the center of mass of the correlations over layers for electrodes in each cluster (Fig. 3b). This indicates that certain electrodes were more similar in their musical encoding to early layers of the transformer model, while others were more similar to later layers. Two of the clusters, cluster 1 and cluster 2, display increasing correlations over layers, but cluster 1 reaches a plateau earlier, while cluster 2 continues increasing in model similarity over layers. Cluster 2 is also located significantly further from posteromedial Heschl’s Gyrus (HG), or primary auditory cortex (Fig. 3c) (*46–48*). We grouped electrodes into anatomical regions (Fig. 3d) which contained at least 5 electrodes each and plotted them on the brain (Fig. 3d and 3e). The electrode clusters display notable anatomical correspondence, with cluster 0 mostly comprising neural sites in primary auditory cortex, cluster 1 including mostly auditory cortex, posterior STG, and planum temporale, and cluster 2 electrodes mostly existing outside auditory cortex and into MTG, temporal pole (T. pole), and frontal gyrus. Nonetheless, electrodes from both clusters 1 and 2 were spread across the entire neural processing hierarchy, demonstrating that contextual processing of music also occurs at several stages of the hierarchy.

**Fig. 3.**
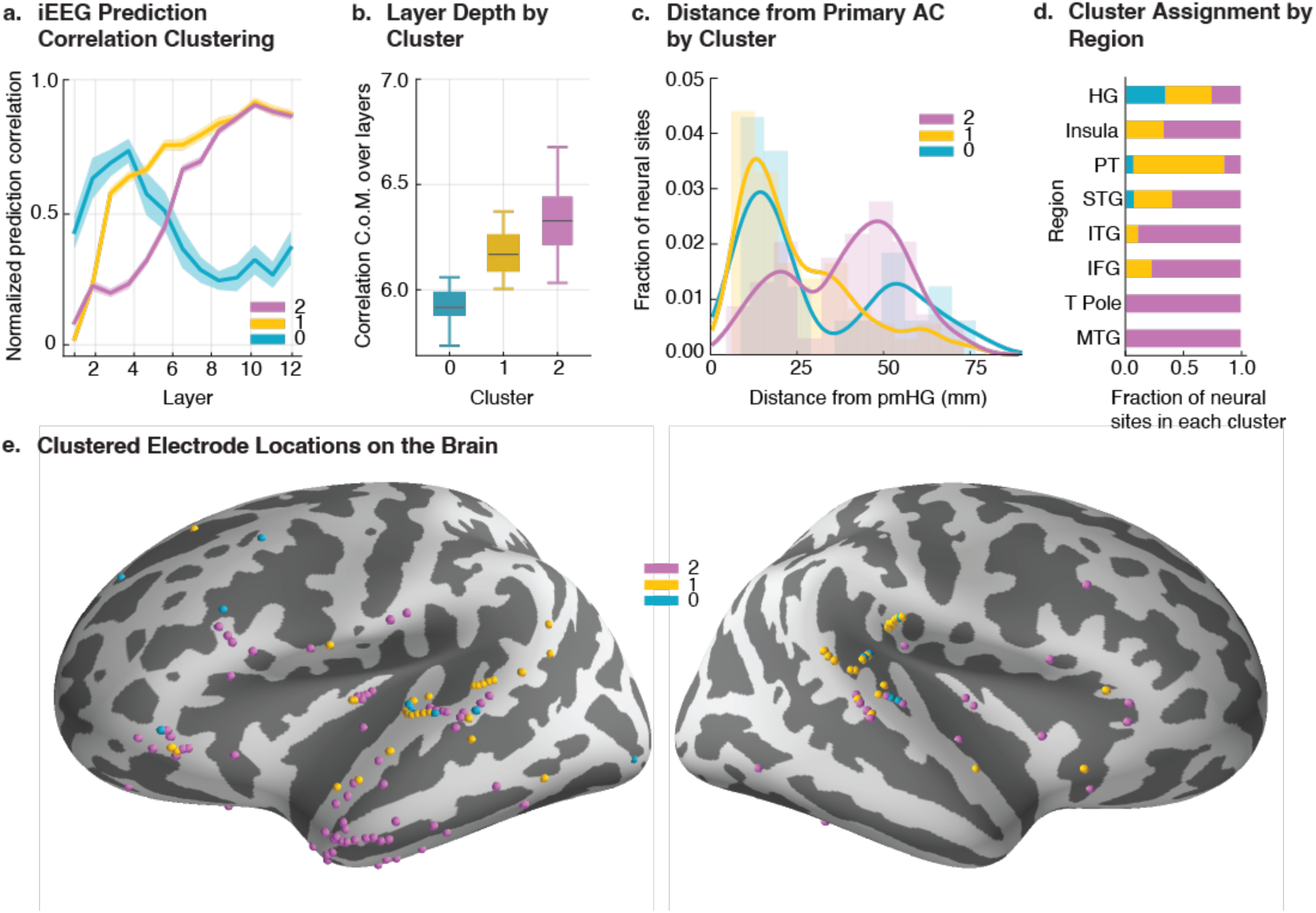
Layer-wise analysis of the prediction correlations of iEEG recordings from the model. **a)** Prediction correlations for each electrode over model layers were normalized between 0 and 1 and clustered into 3 clusters. The cluster averages are plotted over model layers, with shaded regions indicating standard error of the mean over electrodes. **b)** The center of mass (C.o.M.) of the normalized correlations over layers was computed for each electrode and electrodes were grouped by cluster. The distributions of each cluster are shown with box-and-whisker plots. **c)** The distributions of the distances of each electrode from pmHG from each cluster’s electrodes are shown. **d)** Regions of interest were selected based on the presence of at least 5 electrodes, and the fraction of electrodes in each region which were assigned to each cluster are shown. **e)** iEEG electrodes are plotted on an inflated FreeSurfer average brain, with electrode colors indicating cluster assignment.

We then conducted the same analysis as with the EEG recordings by varying the input context window of the transformer embeddings and predicting the iEEG data. The peak score (over layers) and peak scoring layer for the electrodes are shown for each context window (Fig. 4a). On average, the peak score rises and plateaus around 50 notes of context, in agreement with the non-musician EEG results. To investigate the anatomical influence on contextual processing of music in the iEEG data, we quantified the amount of context used by a given electrode using the center of mass of the prediction correlation over context lengths, with a higher value indicating a later rise or plateau in score. This metric for contextual encoding is significantly correlated with the distance of an electrode from primary auditory cortex (Fig. 4b) (Pearson r = 0.32, p = 1.5 × 10^−5^). Thus, electrodes further along the processing pathway utilize more contextual information from the music in their responses. Splitting electrodes into all regions of interest with at least five electrodes each, we found that later regions in the traditional auditory pathway encode context more strongly than earlier regions, such as superior temporal gyrus (STG) encoding more context than HG (Wilcoxon rank sum test, p = 0.028) (Fig. 4c). Interestingly, temporal pole and MTG displayed the strongest contextual encoding (T. Pole p < 0.05 compared to every region except MTG, and MTG with the highest average center of mass but only significant compared to HG, p = 0.029). This measure of context length used by each electrode can also be seen visually over the neural hierarchy in figure 4d. Overall, these results suggest that a hierarchy of musical feature extraction exists in the brain, with greater contextual encoding of music pieces occurring primarily beyond the boundaries of the auditory cortex.

**Fig. 4.**
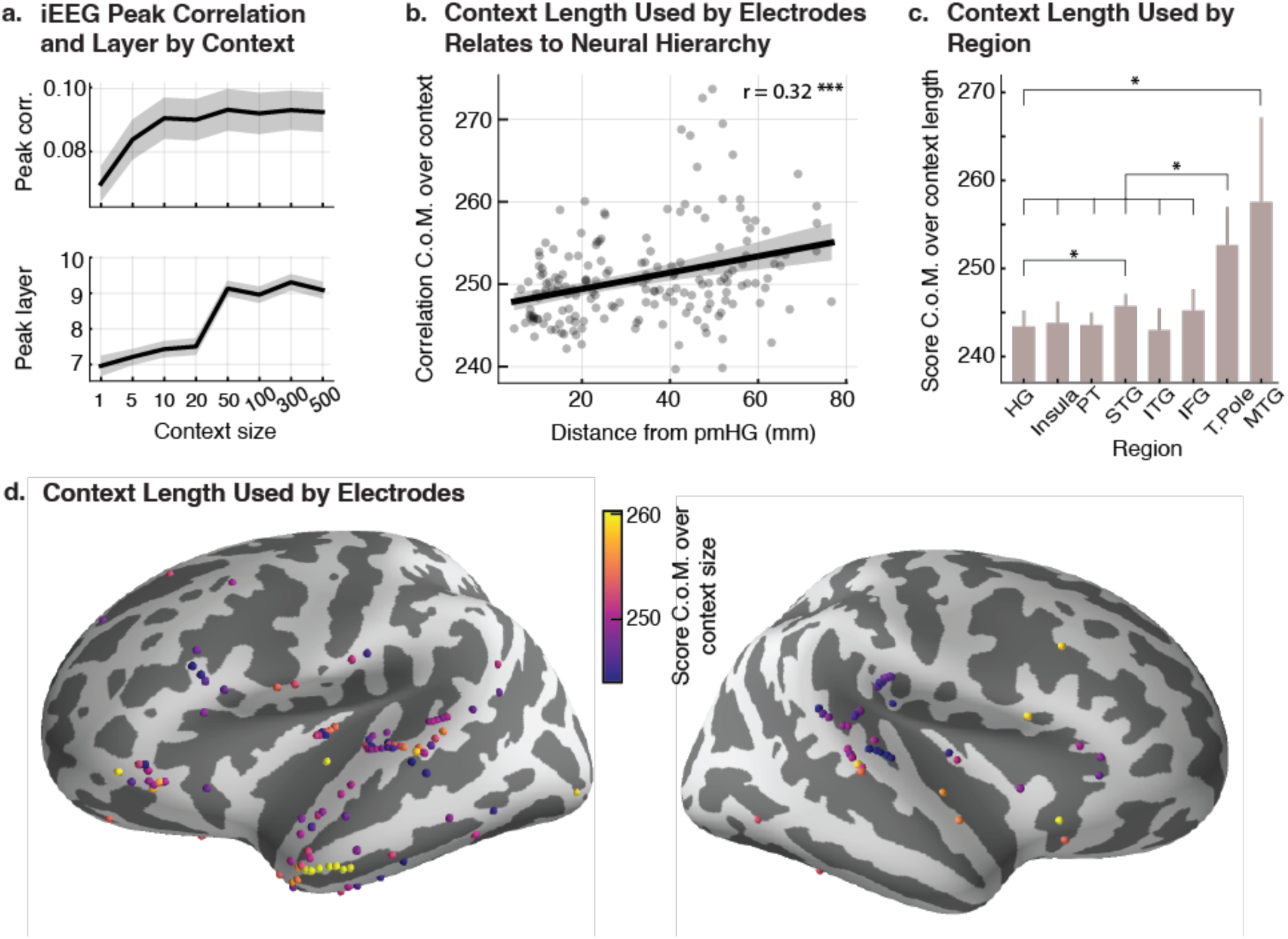
Results from analyzing the relationship between context size and prediction correlations of iEEG recordings. **a)** Peak correlation (over layers) and peak layer of the iEEG electrode prediction correlations when the embeddings were generated from context-limited input. The line shows the average over all electrodes, and the shaded region indicates standard error of the mean. **b)** The center of mass (C.o.M.) of each electrode’s correlations over context sizes was computed and is correlated with each electrode’s distance from pmHG (Pearson r=0.32, p=1.5x10^-5^). Line shows the line of best fit, with the shaded region showing a bootstrapped (n=1000) confidence interval of the line, and *** indicates p<0.001 from the Pearson correlation’s significance test. **c)** The average context size C.o.M. of electrodes in each region of interest is shown, with vertical lines representing standard error of the mean. Statistically significant differences in C.o.M. are shown with the black lines above the bars, where * indicates p<0.05 (Wilcoxon rank sum test). **d)** The context size C.o.M. of each electrode is indicated with its color on the inflated FreeSurfer average brain, clipping the color scale at the 5th and 95th percentiles of the distribution of C.o.M. values.

## Discussion

We studied the embeddings of a music generative transformer and its relation to the human brain’s music processing abilities. We found that the layers of the model demonstrate an increasing processing of musical structure and context, with deeper layers encoding more disentangled features and integrating over longer context, a result which matches similar findings in transformer models trained on text (*42*, *49*). When predicting the EEG responses of musicians and non-musicians from the layers of the transformer model, the neural prediction correlations were greater for musicians compared to non-musicians, especially in later layers and for longer context windows, indicating that musical training may refine the brain’s capacity to construct disentangled representations of music and process musical context. Although prior works has demonstrated stronger baseline encoding in the right hemisphere of melodic content (*50*) and musical syntax (*51*), several previous studies have identified increased left hemisphere recruitment by musicians compared to non-musicians during music note and low-level musical feature processing (*30–32*). Our findings build on these lateralization effects by showing stronger correspondence with disentangled and contextual music features in the left hemisphere by musicians compared to non-musicians. Moreover, our results suggest a distinction in where structure and context are processed in musicians, where the encoding of longer context increasingly involves the frontal cortical areas. To uncover the neural regions behind this contextual processing of music, our iEEG recordings offered more insight. The neural sites which were most similar to the latest layers of the model were distributed throughout the entire auditory processing hierarchy, though they were most prevalent further from primary auditory cortex. Our results build on recent investigations of music feature encoding and contextual integration and suggest that the brain’s processing of musical context is hierarchical, adaptable, and is influenced by cognitive factors such as experience and exposure.

Our analyses suggest that music processing in the brain is hierarchical in nature, especially in the context of varying temporal scales (*22*, *23*). The distinct patterns of neural response to music in musicians and non-musicians when compared to the later layers of the transformer model suggest that higher-order cognitive processes are significantly involved in music listening. This is consistent with previous research that has identified a hierarchical organization in the brain for processing musical elements, ranging from simple features like pitch and rhythm to more complex aspects such as melody and harmony (*22*, *23*, *52*, *53*) as well as long-term syntactic structure (*54*). Additionally, prior research has highlighted the critical role of the frontal cortex, particularly the inferior frontal gyrus, in the hierarchical processing of music (*24–26*, *55*, *56*).

Consistent with these findings, our results demonstrate increased recruitment of the left frontal hemisphere in musicians and stronger correspondence between neural activity in regions beyond the primary auditory cortex and the deeper layers of the model. The progression of predictability in neural responses from the mid to the deeper layers of the transformer in our study may reflect this same hierarchical processing, where basic musical elements are processed at lower levels, and more complex, temporally extended structures are understood at higher levels. Such a multi-tiered processing mechanism is supported by studies showing that different brain regions are activated in response to different musical structures, indicating a distributed yet organized processing network (*57*, *58*). A major implication of our study is that this hierarchical processing may be fine-tuned in musicians, enabling them to more effectively integrate and interpret complex musical structures over time (*59*). A limitation of our methodology is that musicians may have greater familiarity with the Bach pieces used in the study, potentially enhancing their ability to predict upcoming musical structures. This familiarity could partly explain the more disentangled and contextual neural responses observed in musicians, as well as the stronger correspondence between their neural activity and the deeper layers of the model. Consequently, the differences in frontal cortex EEG predictability between musicians and non-musicians may reflect differential recruitment of these regions for generating musical predictions and expectation during listening (*56*). While it remains unclear whether the observed differences are driven primarily by musical training or familiarity, the consistent observation of enhanced neural encoding in musicians highlights the unique ways their brains process music

The differential neural responses to musical context between musicians and non-musicians observed in our study also aligns with the growing body of research indicating that musical training can induce long-lasting functional and structural changes in the brain. For instance, studies have shown that musicians typically exhibit enhanced auditory processing, more robust neural encoding of sound, and greater connectivity in brain regions associated with auditory and motor functions and working memory (*60*, *61*). These adaptations are thought to result from the demands of musical training, which requires precise auditory discrimination and motor coordination, and they manifest as heightened sensitivity and efficiency in processing musical information (*59*, *62*, *63*). Our findings contribute to this understanding by suggesting that such neuroplastic adaptations extend to the processing of musical structure and context, with musicians exhibiting more refined neural encoding patterns, especially in higher-order cognitive processes. This adds a new dimension to our understanding of how musical training shapes the brain, indicating that it not only enhances basic auditory skills but also influences the hierarchical processing of complex musical structures. Future research should delve deeper into these neuroplastic changes, examining how various forms of musical training and exposure distinctively shape the brain’s response to music (*16*, *64*). It would be particularly insightful to explore whether enhanced contextual processing in musicians stems from their established superior working memory (*65*), and to investigate if the underlying mechanisms are akin to those involved in the improved perception of speech in noise, a contextual skill also found to be better developed in musicians (*62*, *66*).

The utilization of advanced AI models, such as the music generative transformer in our study, offers a novel lens through which we can view and understand complex cognitive processes. The success of this model in predicting neural responses to musical stimuli underscores its potential as a proxy for human cognitive processing. This aligns with emerging research suggesting that AI, particularly deep learning models, can simulate aspects of human cognition, offering insights into underlying neural mechanisms (*67*, *68*). In music listening, recent studies have found recurrent and transformer model features to relate to melodic expectations (*27*, *34*). In visual processing, deep learning models like convolutional neural networks (CNNs) have provided insights into the brain’s hierarchical processing of visual stimuli, mirroring the organization of the visual cortex (*69*, *70*). Similar insights have been gained in both recurrent and transformer models of speech and language processing (*71–79*). Our analysis of the contextual encoding of music follows recent findings in language processing which have identified greater contextual dependence in later regions of the neural hierarchy (*77–79*). Several studies have also suggested similarities between linguistic and musical processing through shared neural mechanisms (*24*, *25*, *55*, *56*), providing further evidence that similar transformer architectures can be useful in studying the neural encoding of both types of stimuli. Our study extends the growing field of AI models in neuroscience by demonstrating how a music generative transformer model can simulate the hierarchical and temporal aspects of musical context processing in the brain. We found that such a model revealed distinct differences in this processing between musicians and non-musicians, thereby contributing a novel perspective to the understanding of how musical expertise shapes cognitive functions and neural network models’ role in unraveling these intricate processes. In conclusion, this study advances our understanding of neural processing of music, revealing not just the influence of musical training on the brain, but also the intricate, hierarchical nature of how music is encoded.

## Materials and Methods

### Bach Piano Piece Stimuli

Ten musical pieces from Bach’s monodic instrumental repertoire were converted into monophonic MIDI files and segmented into 150-second snippets. The chosen melodies were originally derived from violin (partita BWV 1001, presto; BWV 1002, allemande; BWV 1004, allemande and gigue; BWV 1006, loure and gavotte) and flute (partita BWV 1013, allemande, corrente, sarabande, and bourrée angloise) scores, and synthesized using piano sounds with MuseScore software (MuseScore BVBA), maintaining a constant tempo (between 47 and 140 bpm). This approach was intended to reduce the familiarity of the pieces for expert pianists while leveraging their preferred instrument timbre to enhance neural responses (*80*). Each 150-second snippet, representing an EEG/ECoG trial, was presented three times during the experiment, resulting in a total of 30 trials presented in a randomized order.

### Calculation of Musical Features

Musical features were derived from the MIDI representation of the musical pieces and included the pitch class, note duration, music piece identity, and full pitch name. Pitch class grouped each note into its respective chromatic category (e.g., C, C#, D), representing the pitch irrespective of octave. Note duration was calculated as the temporal length of each note and normalized to account for tempo variability across the pieces. Music piece identity labeled each note with its corresponding musical piece based on metadata from the MIDI files. Full pitch name combined the pitch class with octave information to assign a unique identifier (e.g., C4, G#3) for each note. These features were analyzed to assess their separability within the embeddings generated by the transformer model and to compare their representations across layers.

### Music Transformer and Temporal Response Function

We used Musicautobot (*37*), a public pre-trained joint encoder-decoder multi-tasking music transformer model, to extract the contextual music embeddings from 8 MIDI files. The music transformer model was trained on four tasks: one for the encoder only, one for the decoder only, and two for the joint encoder and decoder. The encoder was trained on the bidirectional masked prediction objective, and the decoder was trained on the next token prediction task. They were jointly trained to do chord predictions and melody predictions from melody and chord inputs, respectively. The data used to train the model contained classical music from a wide variety of composers, as well as other genres. In this study, we only use the encoder as its bidirectional masked prediction objective is known to preserve the most contextual information. Since the encoder is bidirectional and does not produce causal representations, we masked future note inputs when computing each note’s representations to align with the causal processing of the brain. To perform dimensionality reduction with t-SNE for visualization of a layer’s embeddings, we used the implementation from scikit-learn (*81*) with the default parameters (perplexity = 30, Euclidean distance).

We applied non-negative matrix factorization (NMF) with scikit-learn to reduce the dimensionality of the representations from 2048 to 100, fitting a separate NMF transformation for each layer. We then filled the time-steps between the notes to align these note-aligned transformer features with the EEG or iEEG signals so they would have the same sampling rate.

We trained linear regression models to estimate the temporal response function (TRF) using the mTRF Toolbox (*41*) with L2 regularization to predict the EEG and iEEG responses from the transformer representations. We applied leave-one-out cross-validation (LOOCV) with the regularization term ranging from 10^−6^ to 10^6^ with an increment of power of 10 in the log scale. We used a time-lag window of −300 to 750 ms to fit the TRF models. We employed leave-one-out cross-validation for model fitting, where each model was fitted repeatedly with 9 music pieces and tested on the left-out piece. The average correlation between the predicted neural responses and actual EEG or iEEG signals over these cross-validation folds was reported as an electrode’s correlation.

### Scalp EEG Recording and Preprocessing

Twenty subjects (10 male, aged between 23 and 42, M = 29) participated in the EEG experiment (*13*, *82*). Ten of them were highly trained musicians with a degree in music and at least ten years of experience, while the other participants had no musical background. All subjects provided written informed consent and were paid for their participation. The study protocol was approved by the CERES committee of Paris Descartes University and was performed in accordance with the Declaration of Helsinki. The EEG signals were recorded from 64 electrode positions, digitized at 512 Hz using a BioSemi Active Two system. Audio stimuli were presented at a sampling rate of 44,100 Hz using headphones. Subjects were instructed to maintain visual fixation on a crosshair centered on the screen to minimize motor activities while music was played. We applied a bandpass filter on the normalized EEG signal between 0.5 and 8 Hz using a Butterworth zero-phase filter. The average of the two mastoid channels was used as the reference. The tasks were repeated three times, so we obtained three recordings for each subject and music piece. We fitted a TRF model with all three repetitions combined for each subject and music piece. A non-parametric shuffling paradigm was used to test for electrode significance by shuffling notes 100 times and retraining TRF models for each shuffled set of stimuli. All electrodes for except for occipital electrodes had significant TRF prediction correlations for all subjects (p < 0.05).

### Intracranial EEG Recording and Preprocessing

We recorded iEEG data from six subjects who were undergoing clinical monitoring for epilepsy. The iEEG electrodes were implanted according to clinical need for identifying epileptogenic foci, and any electrodes showing signs of epileptiform discharges, as identified by an epileptologist, were excluded. Prior to electrode implantation, all subjects gave written informed consent for participation in research. The protocol was approved by the institutional review board of North Shore University Hospital. The recordings were preprocessed by first extracting the envelope of the high-gamma band (70-150 Hz) using the Hilbert transform (*83*). This signal has been shown to be highly correlated with neuronal firing rates (*84*, *85*). Each electrode was then downsampled to 100 Hz and used as the neural response target for the TRF models in the same way as for EEG responses. Electrodes were only retained for analysis if they met two significance criteria. The first was a test for acoustic responsiveness using a paired t-test between responses in the half-second before and the half-second after acoustic onsets (p < 0.05, FDR corrected (*86*)). The second was a test for TRF encoding significance, requiring at least 4 out of the 13 model layers to be significantly predictive of the electrode’s response, as judged by the p-values of the trained TRF models (p < 0.05). These p-values were computed from a non-parametric shuffling paradigm, whereby notes were shuffled 100 times and TRF models were trained for each shuffled case, producing the null distribution of correlations for each electrode. All experimental procedures were approved by the local Institutional Review Board (IRB).

### Intracranial EEG Electrode Localization and Code

Each subject’s electrode locations were mapped to the subject’s individual brain by performing co-registration between the pre-implant and post-implant MRI scans with iELVis (*87*). Then, each subject’s electrodes were mapped to the FreeSurfer average brain (*88*) for plotting and inter-subject comparisons. Distance from primary auditory cortex was defined as the Euclidean distance of an electrode from posteromedial HG (TE1.1) (*46*) in FreeSurfer average brain space, since TE1.1 is a common landmark of the beginning of primary auditory cortex (*47*, *48*, *79*). For plotting purposes, all subdural electrodes were snapped to the nearest surface point and plotted on the inflated brain. iEEG neural recordings were preprocessed, including high-gamma envelope extraction, with naplib-python (*89*). TRF models were trained with mTRF toolbox (*41*), and all other analyses were performed with custom scripts in Matlab and Python.

## Data and Materials Availability

The iEEG recordings cannot be made publicly available due to ethical restrictions aimed at protecting the privacy and confidentiality of the human subjects involved in this study. However, the stimuli and the EEG recordings are publicly accessible.

## Acknowledgments

This work was funded by the National Institutes of Health, the National Institute on Deafness and Other Communication Disorders, and the National Science Foundation Graduate Research Fellowship Program. The funders had no influence on study design, data collection and analysis, decision to publish, or preparation of the manuscript.

## Author Contributions

Conceptualization: GM, YAL, NM

Methodology: GM, YAL, SB, ADM, NM

Investigation: GM, YAL, NM

Supervision: NM

Writing – original draft: YAL, NM

Writing – review & editing: GM, NM

